# SPINK3-sperm interaction determines a stable sperm subpopulation with intact CatSper channel

**DOI:** 10.1101/2025.08.26.672324

**Authors:** Anabella R. Nicolli, Xiaofang Huang, Lucia Zalazar, Cintia Stival, Lucila R. Gomez-Olivieri, Gianluca Demare, Ana Romarowski, Mariano G. Buffone, Dario Krapf, Jean-Ju Chung, Andreina Cesari

**Author notes:** Equal contribution (Anabella R. Nicolli, Xiaofang Huang). Corresponding author: Andreina Cesari; Funes 3250 3rd floor, Mar del Plata, Argentina.

## Abstract

Sperm capacitation involves proteolytic remodeling of membrane proteins, including components of the CatSper calcium channel, which is essential for hyperactivation and male fertility. Here, we identify the seminal protease inhibitor SPINK3, a known decapacitation factor that suppresses premature capacitation in the female tract, as the first physiological inhibitor of CATSPER1 processing. In mouse sperm, SPINK3 blocks capacitation-induced CATSPER1 cleavage, preserving a subpopulation with intact CatSper channels and lacking pTyr development in the flagellum. SPINK3 localizes to the outer surface of the sperm principal piece membrane in a CatSper-dependent but non-quadrilateral pattern, stabilizes membrane organization, and delays cholesterol efflux. These results reveal SPINK3 as a multifunctional regulator of capacitation, shaping sperm subpopulations in the female reproductive tract.

**SIGNIFICANCE STATEMENT:** This study unveils a novel physiological regulator from the male reproductive tract, SPINK3, which extracellularly controls the processing of CATSPER1, a key subunit of the CatSper calcium channel essential for sperm capacitation in mice. We show that SPINK3 binds to the sperm surface and prevents the processing of CATSPER1 during capacitation. SPINK3-sperm interaction is dependent on the presence of the sperm-specific CatSper channel. Our findings reveal that SPINK3 stabilizes sperm membranes providing new insights into the molecular regulation of sperm function and fertility.

**GRAPHICAL ABSTRACT:** 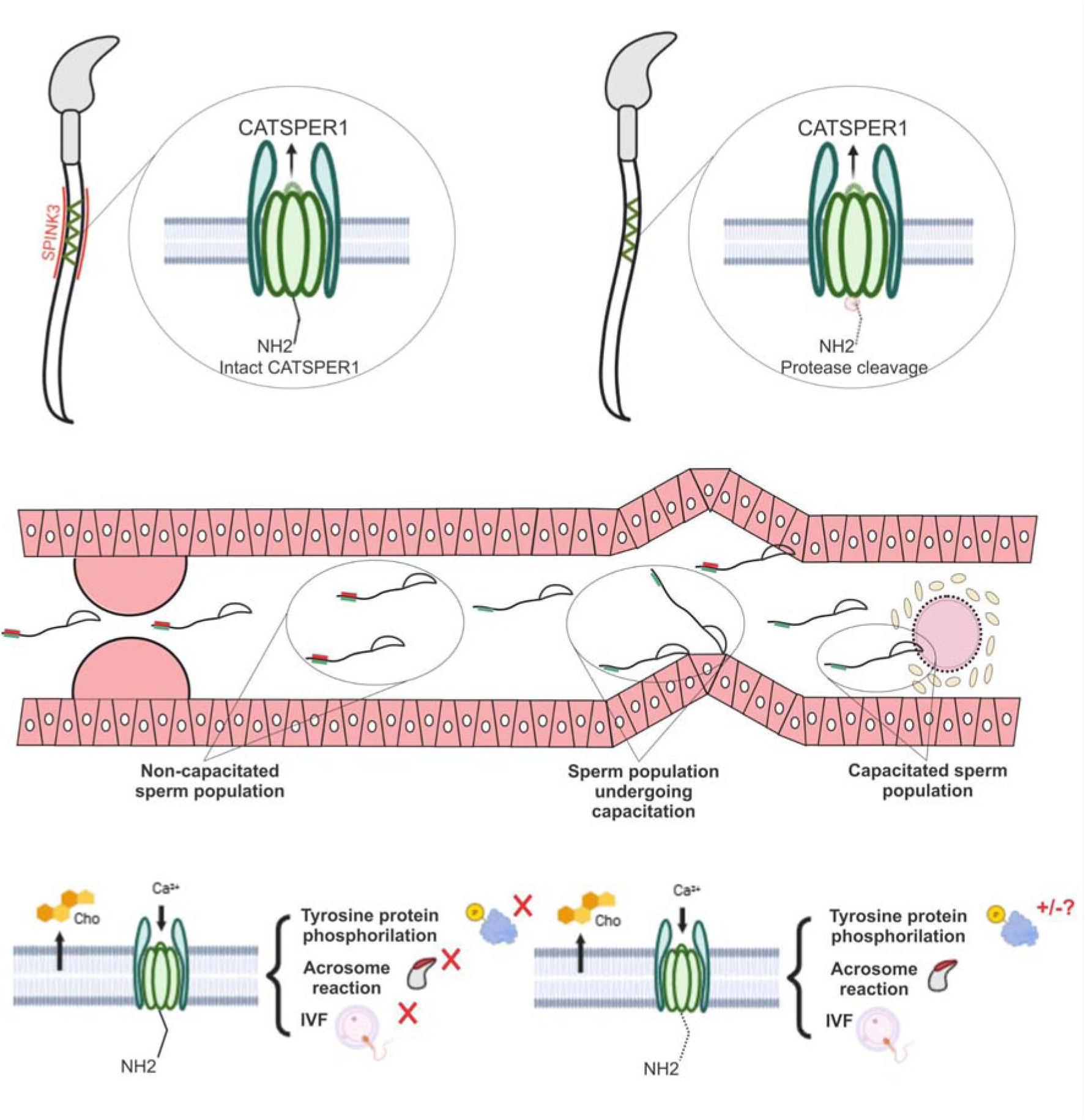

## INTRODUCTION

The sperm journey begins in the male reproductive tract, where millions of sperm undergo surface modifications induced by male secretory molecules (A. RNicolli & Cesari, 2023). After ejaculation, mammalian sperm must undergo biochemical and functional changes within the female reproductive tract, a process collectively known as capacitation (Austin, 1952; Chang, 1951). This process involves membrane remodeling and the activation of intracellular signaling pathways leading to the acquisition of fertilizing ability. Decapacitation factors, typically originating from the male reproductive tract or seminal plasma, coat and stabilize the sperm surface after ejaculation, preventing or delaying capacitation until the sperm reaches an appropriate environment in the female reproductive tract (Dukelow et al., 1967; Leahy & Gadella, 2011; Nicolli & Cesari, 2023).

CatSper is the primary Ca^2+^ channel in mammalian spermatozoa, consisting of four pore-forming subunits (CATSPER1, 2, 3, 4) and at least ten additional auxiliary subunits (Hwang & Chung, 2023). This channel is specifically located in the flagellar membrane of the principal piece, exhibiting pH sensitivity and weak voltage dependence (Kirichok et al., 2006; Ren et al., 2001) and is essential for the development of hyperactivated motility of sperm (Qi et al., 2007; Quill et al., 2003). Infertility has been observed in male mice lacking the *Catsper* genes encoding the transmembrane subunits (Chung et al., 2011; Huang et al., 2023; Hwang, 2024; Qi et al., 2007; Quill et al., 2003; Ren et al., 2001). Capacitation induces specific cleavage and processing of CATSPER1, one of the pore-forming subunits, in the cytosolic N-terminus by unknown sperm enzyme(s), resulting in a graded distribution of heterogeneous sperm populations with varying degrees of CATSPER1 integrity along the female reproductive tract (Chung et al., 2014; Ded et al., 2020). Spermatozoa that reach the ampulla, where fertilization takes place, mostly have intact CATSPER1 signal compared to those that remain behind the utero-tubal junction (UTJ). Recently, it was reported that another CatSper transmembrane subunit, CATSPERθ, which is essential for the CatSper channel assembly and serves as a checkpoint for flagellar trafficking during spermiogenesis, is also processed during sperm capacitation (Huang et al., 2023). This highlights the proteolytic regulation of the CatSper channels as a sperm selection mechanism. Although the cell-permeable potent proteasome inhibitor MG-132, as well as calpain inhibitors and the phosphatase 1/2A inhibitor calyculin A, have been shown to inhibit CATSPER1 processing (Chung et al., 2014; Ded et al., 2020), physiological inhibitors responsible for modulating the processing of CatSper subunits remains unidentified.

SPINK3 (also known as P12 or SPINK1) is a small and highly conserved protein consisting of 80 amino acids, naturally secreted by the male seminal vesicles of most mammals (Dematteis et al., n.d.; Tiwari et al., 2020; Zalazar et al., 2020). It contains a Kazal domain that acts as a substrate analogue by competitively and stoichiometrically binding to the active site of its target protease, forming a stable protease–protease inhibitor complex, such as kallikreins, trypsin and chymotrypsin-like proteases (Assis et al., 2013). SPINK3 also has a sperm binding domain (Lin et al., 2006) and attaches to sperm during ejaculation (Nicolli et al., 2022). SPINK3 is a known decapacitation factor regulating SRC tyrosine kinase activation and membrane potential, apparently without affecting the tyrosine kinase or PKA activity (Zalazar et al., 2020). The BSA-induced increase in intracellular Ca^2+^ concentration ([Ca^2+^]_i_) was demonstrated to depend on CatSper channels. This [Ca^2+^]_i_ increase and CatSper activity are inhibited by the presence of SPINK3 (Zalazar et al., 2020), suggesting an either direct or indirect blockade of calcium entry through the CatSper channel. Similar to other decapacitation factors (Li et al., 2018; Lin et al., 2008; Silva et al., 2021), SPINK3 has been demonstrated to bind sperm during ejaculation until detachment in the female tract, following a specific but asynchronous spatiotemporal kinetics, and contributing to sperm heterogeneity (Nicolli et al., 2022). The protease TESP1 is the only natural target reported for SPINK3 on the sperm head and this interaction was found to be blocked by the soybean trypsin inhibitor, SBTI (Ramachandran et al., 2021).

In this study, we show that SPINK3 regulates CATSPER1 processing and contributes to the generation of sperm subpopulations with different capacitation states. This action is accompanied by the stabilization of the sperm plasma membrane, suggesting that its regulatory effect may involve protease inhibition and membrane-stabilizing mechanisms that prevent protease access to CATSPER1 cleavage sites. These findings highlight a multifaceted role for SPINK3 in modulating sperm capacitation.

## MATERIALS AND METHODS

### Animals

Female and male mice (*Mus musculus*) of BALB/c and C57BL/6 strains and sexually mature (BALB/c x C57BL/6) F1 hybrids were maintained at 22°C with a photoperiod of 12 h light:12 h darkness, food and water *ad libitum*. Sexually mature (2–3 months) male and female mice were euthanized by cervical dislocation.

*Catsper1*-null mice were previously generated (Ren et al., 2001) and maintained on a C57BL/6 background.

Mice were cared for in compliance with the guidelines approved by local Animal Care and Use Committees (UNMdP RD 104/22), following the National Institutes of Health Guide to the Care and Use of Laboratory Animals.

### Antibodies and reagents

Rabbit polyclonal antibodies against CATSPER1 (Ren et al., 2001), CATSPERθ (Huang et al., 2023) and SPINK3 (Sigma HPA027498) were used. A monoclonal anti-HA antibody was purchased from Cell Signaling Technology (#3724), and a monoclonal anti-tubulin antibody (T6094, 1:5000) was obtained from Sigma-Aldrich. HRP-conjugated goat anti-rabbit IgG and goat anti-mouse IgG were used as secondary antibodies. Recombinant SPINK3 protein was produced in *E. coli* and purified using HiTrap IMAC HP (GE Healthcare Life Sciences).

Additional reagents included: Soybean Trypsin Inhibitor (SBTI) (1 µM), Fluo-3 AM (Invitrogen USA, Molecular Probes, F1242), Pluronic acid (Invitrogen, USA), Laminin (100 µg/ml, BD Biosciences), PNA Alexa Fluor 594 lectin (1 µg/ml, Molecular Probes, L-32459), Bodipy-Cholesterol (15 µM, Cayman, 24618), M540 (2.7 µM, Sigma-Aldrich, 323756), FM4-64FX (100 µM, Invitrogen, F34653), CellBrite Fix 640 (2.5 µM, Biotium, 30089-T), Yo-Pro1 (50 nM, Invitrogen, Y3603), Trypsin (347 USP-u/mg, Biological industries, 4192025), PMSF (Phenylmethylsulfonyl fluoride, 500µM, ICN Biomedicals Inc., 195381) and fluorogenic protease substrates Z-CGGR-AMC (7-Amino-4-methylcoumarin), Suc-LLVY-AMC, and Z-SLY-AMC (2=µM each; all from Sigma-Aldrich, Merck, Darmstadt, Germany) .

### Expression and purification of recombinant SPINK3

For the production of SPINK3, the cDNA encoding the mature SPINK3 from *M. musculus* (NCBI ID: NM 009258.5) was cloned into the pET-24b(+) (Novagen, Madison, WI, USA) expression vector. Overexpression of SPINK3 was performed in *Escherichia coli* Rosetta cells (Novagen) and the recombinant protein was purified to apparent homogeneity as described (Assis et al., 2013, and Supplementary Information S1).

### Sperm preparation

Epididymal spermatozoa from adult male mice were collected by swim-out from caudal epididymis in M2 medium (EMD Millipore, MR-015-D) or HM medium (Modified Krebs Ringer Bicarbonate medium: 20 mM Hepes, 119.3 mM NaCl, 4.7 mM KCl, 1.2 mM MgSO_4_, 5.6 mM glucose, 1.2 mM KH_2_PO_4_, 0.5 mM Na pyruvate and 1.7 mM CaCl_2_; pH 7.4). Collected sperm were washed with fresh HM or PBS.

For capacitation, sperm were incubated in HMB (HM plus 15 mM NaHCO_3_ and 5 mg mL^−1^ BSA) or human tubular fluid HTF medium (EMD Millipore, MR-070-D) at (2-7 x 10^6^ cells mL^−1^) w/o purified SPINK3 at 37°C, 5% CO_2_ for 1 or 2 hr. SPINK3 concentrations (5-13 µM) varied according to the assay.

In a set of experiments capacitation was performed in the presence of SPINK3 (13 µM) or SBTI (1 µM) (Noda & Ikawa, 2019; Samanta et al., 2018). For the preparation of sperm lysates, sperm samples (10 × 10= cells) were resuspended in 1=mL of 0.1% Triton X-100 in PBS. The suspension was sonicated three times for 1 second each using a small ultracentrifuge tube. Samples were then agitated for 1 hour at 4=°C, followed by centrifugation at 15,000=×=g for 10 minutes at 4=°C. The supernatant was collected and used immediately. Lysates were always prepared freshly before use.

### Seminal vesicle lysates

Seminal vesicle homogenate was prepared from seminal vesicles obtained from adult male mice. The tissues were placed in ice-cold PBS (pH 7.4) and homogenized using a IKA T10 basic homogenizer under cold conditions. The homogenate was then centrifuged at 10000 × g for 10 minutes, and the supernatant was filtered (0.22µm) and stored on ice until use. The homogenate was subsequently used in activity assays, adjusting the volume based on the estimated concentration of native SPINK3 in seminal vesicle extracts.

### Proteolytic activity of sperm lysates and inhibitory assay

Proteolytic activity of sperm lysates was assessed over fluorogenic substrates: Z-Arg-Gly-Arg-AMC (for trypsin-like proteases), Suc-LLVY-AMC (for chymotrypsin-like proteases), and Z-SLY-AMC (for elastase-like proteases), each at a final concentration of 2=µM. All reactions were performed in 96-well black plates (Nunc) with a final volume (Fv) of 100=µl per well.

Reaction mixtures were prepared at room temperature with 40 µl of sperm lysate as protease source. The reactions (Fv=100 µl) contained 50 mM Buffer phosphate pH 7.4, w/o the following inhibitors: SPINK3 (1.3=µM final), SBTI (2=µg/ml final), seminal vesicle homogenate as a control of native SPINK3 (6=µl) and PMSF (500 µM). The reaction was initiated by adding 2=µM CGGR-AMC substrate, and fluorescence (Ex 355=nm / Em 436=nm) was measured every minute for 30=min at 37=°C with agitation (Fluoroskan Ascent microplate reader, Thermo Electron Corporation, Waltham, MA, USA). Blank reaction was conducted including 40 µl 0,1% Triton. Enzymatic activity was calculated as Δfluorescence over time and expressed relative to the sperm lysate fluorescence. Inhibition is evidenced as a decrease in proteolytic activity.

### In vivo proteolysis of CATSPER1

Capacitation was performed w/o 13 µM SPINK3 or 1 µM SBTI for 2hrs in HTF media. Western blots with whole sperm lysates (1 or 2 x 10^6^ cells/lane) were performed and developed with anti-CATSPER1 (Ren et al., 2001), anti-CATSPERθ (Huang et al., 2023), anti-HA (CST, #3724) or anti-acetylated Tubulin (Sigma, T7451), followed by the corresponding HRP-conjugated goat anti-rabbit or anti-mouse IgG.

### In vitro proteolysis of CATSPER1

HEK293T cells were transiently transfected with construct encoding CATSPER1-HA with Lipofectamine 2000 (Invitrogen), following the manufacturer’s instruction. CATSPER1-HA was solubilized with 0.1% Triton X-100 PBS (PBST) buffer, containing EDTA-free protease inhibitor cocktail and pulled-down with anti-HA magnetic beads (Pierce, 88836) for 1 h. at RT. The enriched CATSPER1-HA protein beads were incubated with 30μL of sperm lysates solubilized from 3.0×10^5^ sperm cells w/o 13 µM SPINK3 at 37°C for the indicated times. After incubation, the mixture was denatured by LDS sampling buffer supplemented with 50 mM DTT at 75°C for 15 min. Western Blot was performed as described before.

### Time-lapse live cell calcium Imaging

Mouse epididymal spermatozoa (n=3) (10 x 10^6^ cells/ml) were loaded with Fluo-3 AM (10 µM) (Invitrogen USA, Molecular Probes F1242) and 0.02% (v/v) pluronic acid (Invitrogen, USA) as a surfactant at 37°C for 20 minutes in a non-capacitating medium (NC). Non-capacitated sperm were then incubated w/o SPINK3 (13 µM) or SBTI (1 µM) for 15 minutes at 37°C and then, adhered to a fluorescence compatible 96 well plate previously covered with laminin (100 µg/ml, BD Biosciences). HM medium supplemented with 15 mM NaHCO_3_ was added to the chamber and basal fluorescence was monitored under confocal microscope (excitation 488 nm; emission 526 nm, Nikon C1SiR) at 400x magnification. BSA (5 mg/ml) in a minimum volume was applied, and changes in the calcium signal were recorded throughout the process. The viability of the spermatozoa was determined through the motility of the flagellum. The intensity of the fluorescence signal in the head of spermatozoa with motile flagella was quantified using the free software ImageJ 1.43 (National Institutes of Health, Bethesda, Maryland, United States). Data was normalized using the equation (F-F0)/F0, where F is the fluorescence intensity at time t and F0 is the mean fluorescence taken during the period before the addition of BSA (initial 10 seconds). Then, the total series of (F-F0)/F0 was plotted against time, as well as the percentage of positive response to the stimulus.

### Acrosomal reaction (AR) assay

Mouse epididymal spermatozoa (n=3) were capacitated (7.5 x 10^6^ cells/ml) for 60 minutes w/o SPINK3 (13 µM) or SBTI (1 µM). To induce the acrosomal reaction, spermatozoa were incubated with progesterone (50 µM) (Zalazar et al., 2020) or Ca^2+^ ionophore A23187 (10 µM) (Sigma-Aldrich, United States) for 15 min at 37°C after capacitation. The cells were then incubated for 15 minutes at 37°C with the PNA Alexa fluor 594 lectin (1 µg/ml, Molecular Probes, L-32459), which allows AR to be detected by binding to glycoproteins of the internal acrosomal membrane. The fluorescence was observed under a microscope at a magnification of 400x. Sperm that lost their acrosomes (red fluorescence) were expressed as a percentage of 100 cells per replicate.

### Cholesterol efflux assay

Mouse epididymal spermatozoa (n=3) (15 x 10^6^ cells/ml) were loaded with Bodipy-Cholesterol (15 µM) (Cayman, 24618) for 30 minutes at 37°C and incubated 15 minutes at 37°C w/o SPINK3 (13 µM) or SBTI (1 µM). Cells were washed to remove excess reagent and exposed for 60 minutes to either capacitation (CAP) or non-capacitation (NC) conditions. The fluorescence of the labeled Bodipy cholesterol was observed in a Nikon Eclipse T2000 epifluorescence microscope with a magnification of 400x, and the percentage of cells that preserved the labeled cholesterol was quantified.

### Determination of lipid disorder

Mouse epididymal sperm (n=3) (10×10^6^ cells/ml) were incubated in the presence or absence of rSPINK3 (13 µM) for 15 minutes at 37°C in HM medium. Subsequently, they were incubated under CAP or NC conditions. After capacitation, 2.7 µM M540 (Ex 555/Em 578 nm, Sigma-Aldrich, 323756) was added to the sperm suspension (1×10^6^ total cells) and incubated for 15 minutes at room temperature in the dark. Cells were washed and Yo-Pro1 was added at a concentration of 50 nM (Ex 491/Em 509 nm., Invitrogen, Y3603). The samples were analyzed by flow cytometry. The results were analyzed using the open access Flowing 2 software. During acquisition, sperm were discriminated from debris. The results are expressed as the percentage of cells with lipid disorder over the total live cells (M540+/Yo-Pro1-).

### Immunolabeling of SPINK3 and CATSPER1

Epididymal spermatozoa from WT and *Catpser1*-/-mice (n=3) were incubated in non-capacitating medium w/o SPINK3 (13 µM) for 15 minutes at 37°C. Each condition was centrifuged twice at 500 × g for 5 minutes, and re-suspended in the same volume of fresh PBS medium without SPINK3. Cells were seeded on 8-well glass slides and allowed to air-dry. Sperm were fixed with 3.7% paraformaldehyde in PBS for 15 minutes at room temperature, washed with PBS and permeabilized with 0.5% Triton X-100 for 5 minutes. After washing with PBS, cells were blocked with 5% BSA in PBS for 1 h at room temperature and followed by overnight incubation with anti-SPINK3 (1:50 in TBS supplemented with 1 % BSA and 0.1 % tween, Sigma HPA027498) at 4 °C. After washing with PBS, anti-rabbit IgG-Alexa 488 (1:1000, Abcam, ab15105) was added for 2 hours at room temperature. Finally, the cells were washed with PBS and mounted to acquire images with the fluorescence microscope NIKON ECLIPSE E800. Images from both conditions (WT and *Catpser1*-/-) were equally processed and analyzed with ImageJ (v1.38, NIH). For data analysis, the percentage of SPINK3-positive sperm under each condition was adjusted by subtracting the count of false positive sperm observed in the basal immunofluorescence controls (i.e., WT or *Catpser1*-/-sperm incubated without SPINK3).

To assess tyrosine-phosphorylated proteins (anti-pY), sperm were incubated w/o SPINK3 (13 µM), incubated under CAP or NC conditions and then the cells were fixed with 4% formaldehyde, washed, and placed on slides. Once dried, they were fixed with 96% ethanol, permeabilized with 10 µl of T-PBS (1x PBS, 0.5% Tween 20%) with 0.5% Triton 100 for 20 minutes. After washing, they were blocked with 3% BSA in T-PBS for 1 hour. The samples were washed and incubated for 1 hour with the primary antibody anti-tyrosine mouse monoclonal (anti-pY) (1:500, Clone 4G10, Millipore) or anti-SPINK3 (1:50, Sigma Aldrich, HPA027498) in blocking buffer, followed by the secondary antibodies anti-rabbit IgG-Alexa488 (1:1000, Invitrogen A11070) and anti-mouse IgG-Alexa455 (1:500, Invitrogen, A21425). The preparations were mounted with glycerol:PBS (9:1) and observed under a fluorescence microscope with a magnification of 400x (Ex 480/Em 525 nm).

### Structured illumination microscopy

2D structured illumination microscopy (SIM) imaging was performed with Zeiss LSM710 Elyra P1 using alpha Plan-APO 100X/1.46 oil objective lens. Samples were prepared as described in the previous section (Sperm labelling with SPINK3 and immunostaining with CATSPER1 antibody). Raw images were processed and rendered using Zen 2012 SP2 software (Carl Zeiss).

### Plasma membrane labeling

After obtaining the sperm sample and co-incubating it with SPINK3 as previously described, the sample was incubated with 100 µM FM4-64 FX for 3 minutes at room temperature or with 2.5 µM CellBrite Fix 640 for 15 minutes at 37°C, followed by the SPINK3 immunolabeling protocol as described above. Confocal images were acquired using a Zeiss LSM-880 microscope equipped with an Alpha Plan-Apochromat 100x/1.46 objective. Images were also captured in Super-Resolution mode using an Airyscan detector. A total of 10 frames were acquired, and the Mean-Shift Super Resolution (MSSR) algorithm (Torres-García et al., 2022) was applied to enhance resolution.

### In vitro fertilization

F1 hybrid females (female Balb/c x male C57BL/6) were superovulated by intraperitoneal injection with 7.5U of PMSG and 48 hours later with 5U of hCG. Cumulus-oocyte complexes (COCs) were obtained from the ampulla 15 hours post hCG and placed in G-IVF plus medium droplets (LifeCell) under mineral oil (M8410, Sigma-Aldrich). Subsequently, groups of COCs were transferred to a clean medium drop (90 µl) under oil. Spermatozoa were obtained from C57BL/6 males in HM medium, incubated w/o SPINK3 (13 µM) for 15 minutes at 37°C and capacitated in G-IVF plus medium (LifeCell) (5 x 10^6^ cells/ml). COCs were inseminated with 10 µl of the sperm suspension and incubated at 37°C, 5% CO2 for 4 hours. Afterward, COCs were washed, and putative zygotes were incubated for 24 hours and 5 days (post-IVF) to assess the two-cell and blastocyst stages under an inverted microscope with a magnification of 400x (Nikon Eclipse Ti-S). Remanent sperm were prepared for immunostaining. Bar charts show the percentage of two-cell embryos and blastocysts over the total fertilized oocytes.

### Statistical analysis

Data were analyzed by GLMM (generalized linear mixed effect model) to determine statistical significance between treatments and control (Zuur, 2019). The model included the male variable as a random effect. Normality of residuals was assessed by plotting theoretical quantiles vs standardized residuals (Q–Q plots). Homogeneity of variance was evaluated by plotting residuals vs fitted values. A post hoc analysis was conducted with the ‘lsmeans’’ package (Lenth, 2016). In all cases, the analysis was performed using R software version 3.3.33, with the ‘nlme’ package for Gaussian models (Pinheiro et al., 2020). Statistically significant differences were determined at P < 0.05. Graph bars indicate mean ± s.e.

## RESULTS

### SPINK3 inhibits CATSPER1 processing induced by *in vitro* capacitation

It has previously been reported that the cytoplasmic N-terminal domain of the CATSPER1 undergoes proteolytic processing in sperm cells during capacitation, and the progression of this processing is negatively correlated with the appearance of tyrosine phosphorylation at single cell level *in vitro* and *in situ* (Ded et al., 2020). More recently, another CatSper subunit, CATSPERθ, which plays a role in CatSper channel assembly and serves as a checkpoint for CatSper flagellar trafficking, was also found to undergo capacitation-induced degradation (Huang et al., 2023).

SPINK3 has previously been shown to inhibit CatSper channel activity (Zalazar et al., 2020). As this small protein is not only a decapacitation factor but also a protease inhibitor, we hypothesize that SPINK3 might be related to the regulation of CATSPER1 and/or CATSPERθ processing. To test this idea, we evaluated capacitation-induced CATSPER1 and CATSPERθ protein degradation in mouse sperm incubated under capacitating conditions, with or without SPINK3. Interestingly, we found that the presence of SPINK3 abolished the degradation of CATSPER1, but not CATSPERθ (**Figure 1A**). Consistent results were seen with anti-CATSPER1 immunostaining in mouse spermatozoa (**Figure 1B, Supp. Figure 1**). As sperm contains several proteases (Cesari et al., 2010), we assessed the proteolytic activity of sperm lysates using synthetic substrates specific for trypsin, chymotrypsin, and elastase, in order to identify the protease family targeted by SPINK3. Our results confirmed that trypsin-like activity was predominant, and that this activity was inhibited by SPINK3, by seminal vesicle lysates (which contain native SPINK3), and by commercial inhibitors such as PMSF (phenylmethylsulfonyl fluoride) and SBTI (soybean trypsin inhibitor) (**Figure 1C**). As a control, we capacitated mouse spermatozoa in the presence of SBTI and observed the same blocking effect on CATSPER1 processing (**Supp. Figure 2**). Heterologously expressed full-length recombinant CATSPER1 has also been validated for processing by sperm lysate *in vitro* due to the presence of specific endogenous protease(s) (Ded et al., 2020). Similarly, this *in vitro* CATSPER1 processing was also inhibited by the presence of SPINK3 (**Figure 1D**). Taken together, these results show that SPINK3 interferes with proteolytic processing of CATSPER1, by inhibiting sperm trypsin-like protease(s). Furthermore, when sperm were preincubated with recombinant SPINK3 under capacitating conditions, only spermatozoa lacking SPINK3 showed protein tyrosine phosphorylation (pTyr) development in the flagella (**Figure 1E**), consistent with the previously reported inverse correlation of pTyr and CatSper channel integrity (Chung et al., 2014; Ded et al., 2020; Hwang & Chung, 2023). These results support the idea that SPINK3 binding results in a population of sperm with intact CATSPER1 and SPINK3 bound to the flagellar surface.

**Figure 1.**
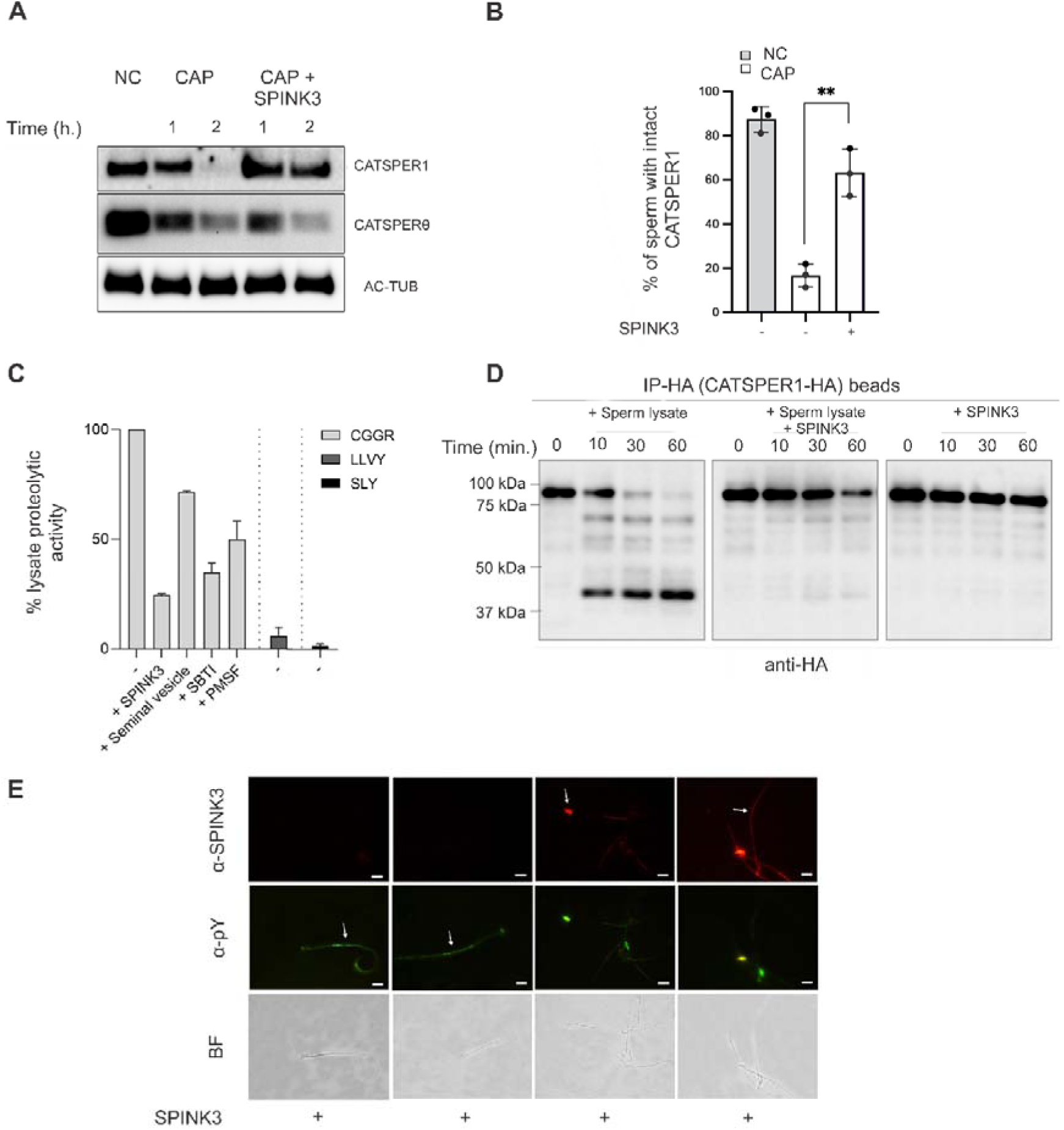
Capacitation-induced CATSPER1 processing by sperm proteases is inhibited by SPINK3 *in vitro*. **A)** Mouse epididymal spermatozoa were incubated w/o SPINK3 (13 µM) and exposed to capacitating conditions for either 1 or 2 hr., as indicated. Immunoblot analysis of non-capacitated (NC) or capacitated (CAP) sperm lysates was performed with anti-CATSPER1 antibody raised against the 1–150 aa region of CATSPER1 (7) and anti-CATSPERθ antibody, raised against the C-terminus of CATSPERθ (12). Acetylated tubulin (AC-TUB) was probed as loading control (n=3). **B)** Immunostaining with anti-CATSPER1 antibody was performed on non-capacitated sperm or capacitated sperm that were previously incubated w/o SPINK3 for 30 min. Sperm with intact CATSPER1 signal were quantified (n=3). **p < 0.001. **C)** Proteolytic activity of sperm lysates was measured over three fluorogenic substrates—Z-Arg-Gly-Arg-AMC (CGGR; trypsin substrate), Suc-LLVY-AMC (LLVY; chymotrypsin substrate), and Z-SLY-AMC (SLY; elastase substrate) in the presence or absence of SPINK3 (13 µM), seminal vesicle homogenate (native SPINK3 source), SBTI (soybean trypsin inhibitor) (2=µg/ml), or PMSF (500µM) (serine protease inhibitor) (n=3). Enzymatic activity was calculated as Δfluorescence over time and expressed relative to the sperm lysate. **D)** FL-CATSPER1-HA was expressed in 293T cells and pulled-down using magnetic beads conjugated with anti-HA antibody. The recombinant protein was then incubated with sperm lysates w/o SPINK3 (13 µM) at 37 ℒC for 0, 10, 30, or 60 min and subjected to immunoblot with anti-HA antibody. **E)** Tyrosine phosphorylated proteins and sperm-bound SPINK3 were detected by immunofluorescence, with anti-pY (clone 4G10, green) and anti-SPINK3 (HPA027498, red) antibodies, respectively. Arrows indicate spermatozoa exclusively stained with anti-pY or anti-SPINK3. Scale bar: 10 μm.

### SPINK3 binds to the sperm flagellar membrane in a CatSper-dependent manner

The CatSper channel forms four linear nanodomains along the flagellar membrane in which the lipid raft marker caveolin1 resides (Chung et al., 2014). Recombinant SPINK3 can bind to raft membrane domains in both acrosomal and tail membranes of non-capacitated mature epididymal spermatozoa and this union is avoided by choleratoxin B supporting binding to GM1 (A. Nicolli et al., n.d.; Zalazar et al., 2020). Given that SPINK3 is a secretory protein, we tested whether SPINK3 and CatSper co-localize in the quadrilinear nanodomains. Confocal microscopy shows that, unlike CATSPER1, which is localized exclusively to the principal piece, SPINK3 is present in both principal piece and end piece of the flagellum (**Figure 2A, upper panel**). The distinct linear CatSper nanodomain arrangement visualized by Structured Illumination Microscopy (SIM) was not seen for SPINK3 in the flagellar membrane (**Figure 2B**). These observations, combined with our finding that SPINK3 inhibits the processing of the N-terminal cytoplasmic domain of CATSPER1, prompted us to further investigate the spatial distribution of SPINK3 in relation to the plasma membrane. Thus, SPINK3 immunostaining was performed together with membrane labelling using the specific plasma membrane probes FM4-64 and CellBrite. Images from Airyscan microscopy and MSSR indicate that SPINK3 is externally associated with the plasma membrane (**Figure 2C, Supp. Movie 1**). Accordingly, SPINK3 signal spans a wider area than the measured membrane-to-membrane distances (**Figure 2C, MSSR Images**). Nevertheless, we observed that the SPINK3 association with sperm is compromised in *Catsper1* KO sperm and there is also a pattern shift (**Figure 2A (lower panel), D; Supp. Figure 3**). It is possible that in *Catsper1* KO sperm the loss of CatSper complex from flagellum changed the membrane property and/or protease localization, thus compromising the attachment of SPINK3. These results suggest that SPINK3 interacts with the flagellar membrane in a CatSper-dependent manner but does not directly bind to CatSper.

**Figure 2.**
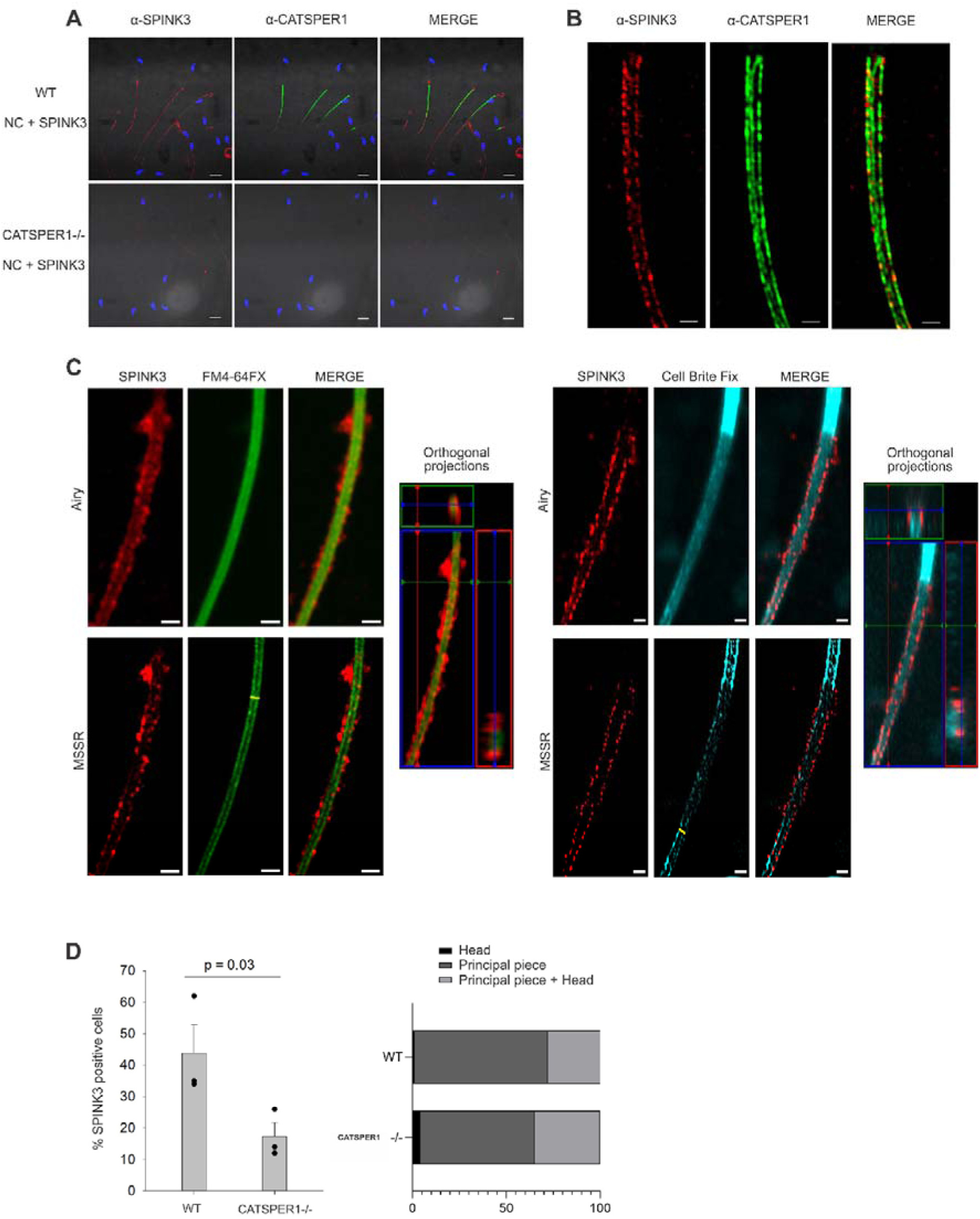
Immunodetection of recombinant SPINK3 binding to epididymal spermatozoa in wild type (WT) and *Catsper1* KO (*Catsper1-/-*) mice. **A)** Non-capacitated (NC) epididymal spermatozoa from WT and *Catsper1*-/-mice (n=3) were preincubated with SPINK3 (5 μM), and immunostaining was performed to detect SPINK3 (red) and CATSPER1 (green). Nuclei were counterstained with Hoechst (blue). Representative images from three independent experiments are shown. Scale bar: 10 µm. **B)** 2D Structured illumination microscopy (SIM) images reveal the plasma membrane localization of SPINK3 (red) on the WT NC sperm cells. The immunostaining of CATSPER1 (green) was used to indicate the boundary of the plasma membrane. Three of the four lines of CatSper nanodomains (as indicated by CATSPER1, green) were seen in this confocal microscopy image. **C)** (Left panel) Airyscan-and MSSR-processed images show SPINK3 (red) localized on the plasma membrane of the sperm principal piece, labeled with FM4-64 FX (green). At the level of the yellow line in the FM4-64 FX MSSR image, the distance measured for FM4-64 FX is 0.409 µm, whereas for SPINK3, it is 0.736 µm. Orthogonal projections from a Z-stack are shown. Scale bar: 1 µm. (Right panel) Airyscan-and MSSR-processed images show SPINK3 (red) on the plasma membrane of the sperm principal piece, labeled with CellBrite Fix (cyan). The midpiece plasma membrane exhibits stronger staining than the principal piece. At the level of the yellow line in the CellBrite FixMSSR image, the distance measured for CellBrite is 0.442 µm, whereas for SPINK3, it is 0.752 µm. Orthogonal projections from a Z-stack are shown. Scale bar: 1 µm. **D)** (Left panel) Quantification of SPINK3-positive cells in WT and *Catsper1*-/-spermatozoa, analyzing both head and tail labeling. Data represent the mean ± SEM (n=3). Significant differences (p < 0.05) between WT and *Catsper1*-/-groups are indicated. (Right panel) Distribution of SPINK3 labeling in spermatozoa head and tail regions, expressed as percentages for WT and *Catsper1*-/-samples.

### SPINK3, but not SBTI, specifically affects sperm capacitation events

Given that protection of CATSPER1 from proteolytic processing (*i.e.*, intact CATSPER-mediated Ca^2+^ signaling) is required to control the timing of capacitation, we analyzed whether the protease inhibitory activity of SPINK3 accounts for its function as a decapacitation factor. To this end, we first tested whether the trypsin inhibitor SBTI could mimic the known effects of SPINK3 on capacitation since SBTI can also inhibit CATSPER1 processing (**Supp. Figure 2**). In contrast to SPINK3, there was no difference in the BSA-induced increase in [Ca^2+^]_i_ between SBTI-treated and untreated spermatozoa as assessed by single-cell Ca^2+^ imaging with Fluo-3 AM (**Figure 3A**), nor in the percentage of acrosome-reacted spermatozoa (**Figure 3B**). Notably, neither SPINK3 nor SBTI affected intrasperm alkalinization (**Figure 3C**), which is known to increase CatSper currents (Ferreira et al., 2021; Hwang et al., 2019; Kirichok et al., 2006). All these results suggest that among trypsin-like protease inhibitors, SPINK3 specifically inhibits capacitation events such as rise in [Ca^2+^]_i_ and AR likely through a mechanism distinct from its inhibition of CATSPER1 cleavage.

**Figure 3.**
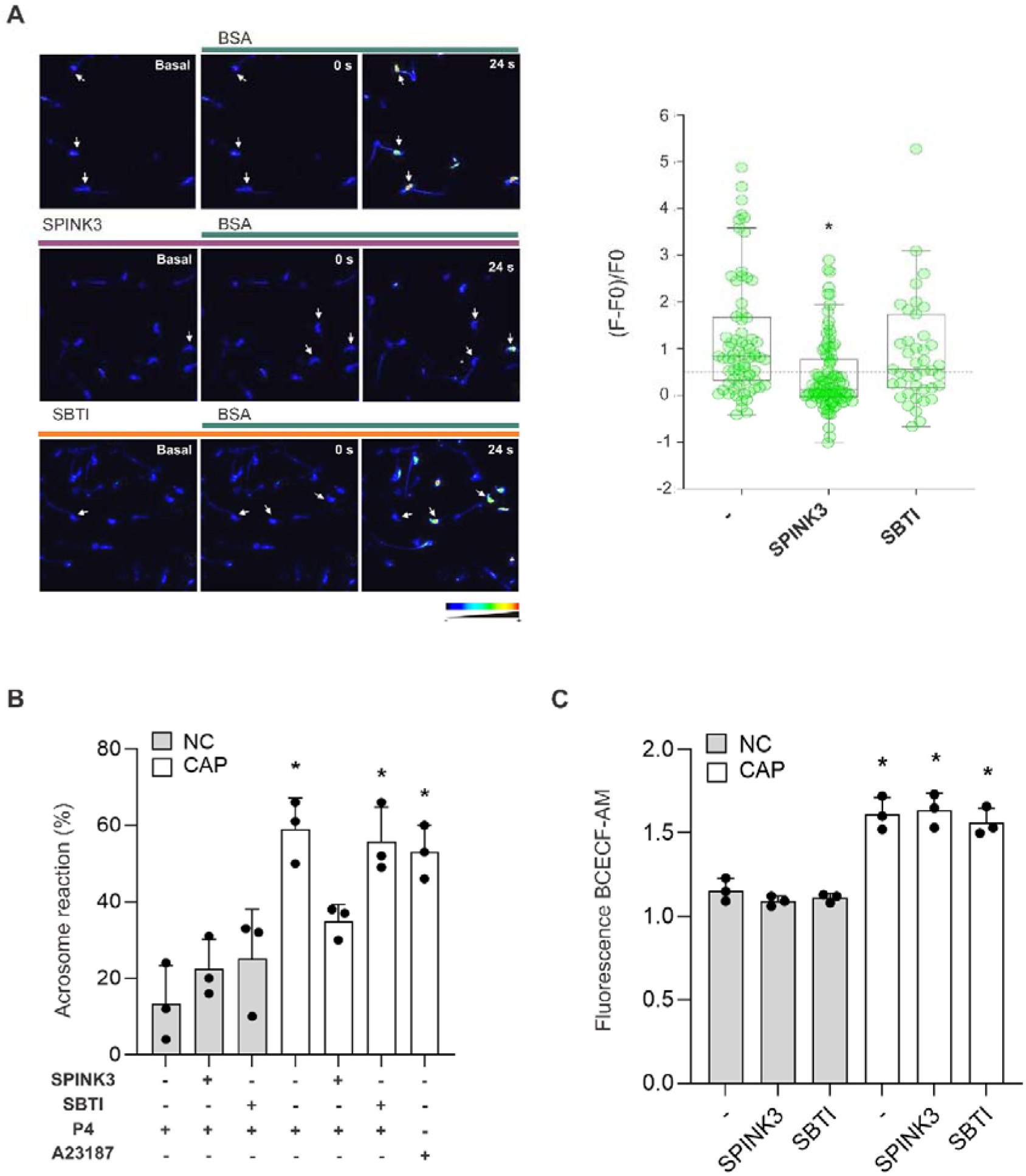
The inhibitory effect of SPINK3 on capacitation is not mimicked by a standard trypsin inhibitor. **A)** Non-capacitated (NC) sperm loaded with Fluo-3 AM were exposed to SPINK3 (13 µM) or SBTI (1 µM). After washing with fresh media containing NaHCO_3_, sperm cells were attached on laminin-coated slides and calcium signal (F0) was monitored before and after BSA was added (left panels). The fluorescence increments of individual live cells (F-F0)/F0 after 24 sec after BSA addition were plotted for each condition: control (-), SPINK3 or SBTI (right panels). *p < 0.05 significant differences in the percentage of responsive cells (>0.5 fold), compared to control untreated sperm. **B)** Acrosome reaction was induced by 50 µM progesterone (P4) after capacitation. As control, NC sperm cells were incubated with the Ca^2+^ ionophore A23187 (10 µM). Data represent mean ± SE; N = 3. *p < 0.05 significant differences compared to NC sperm. **C)** Cells under capacitating (CAP, 60 min) or non-capacitating (NC) conditions were loaded with the pH-sensitive probe BCECF-AM (2 µM), and fluorescence was measured (Ex 520/Em 605 nm). *p < 0.05 significant differences compared to NC sperm.

### SPINK3 alleviates capacitation-induced membrane perturbations

Cholesterol efflux from the sperm plasma membrane is a crucial step in capacitation. These changes promote an increase in membrane fluidity or decrease stability (Flesch et al., 2001) and can be measured using BODIPY-cholesterol (BODIPY/Ch), a fluorescent compound that can replace endogenous cholesterol in the membrane without disturbing the lipid acyl chain region. Therefore, its loss can be measured by changes in fluorescence intensity (Bernecic et al., 2019; Sankaranarayanan et al., 2011). By contrast, the subsequent membrane remodeling can be assessed by the incorporation of anionic lipophilic fluorescent dye Merocyanine 540 (M540) as the amount of dye per unit surface area increases in less-ordered membranes (McEvoy et al., 1988; Steckler et al., 2015). Using these lipid indicators, we show that capacitated spermatozoa contain lower cholesterol content (**Figure 4A**) and increased lipid disorder (**Figure 4B**) compared to non-capacitated sperm, as previously reported. In contrast, pre-incubation with SPINK3 blocked the capacitation-associated cholesterol efflux and lipid disorder (**Figure 4A**, **B**), suggesting that SPINK3 binding to the membrane hinders the membrane reordering during capacitation.

**Figure 4.**
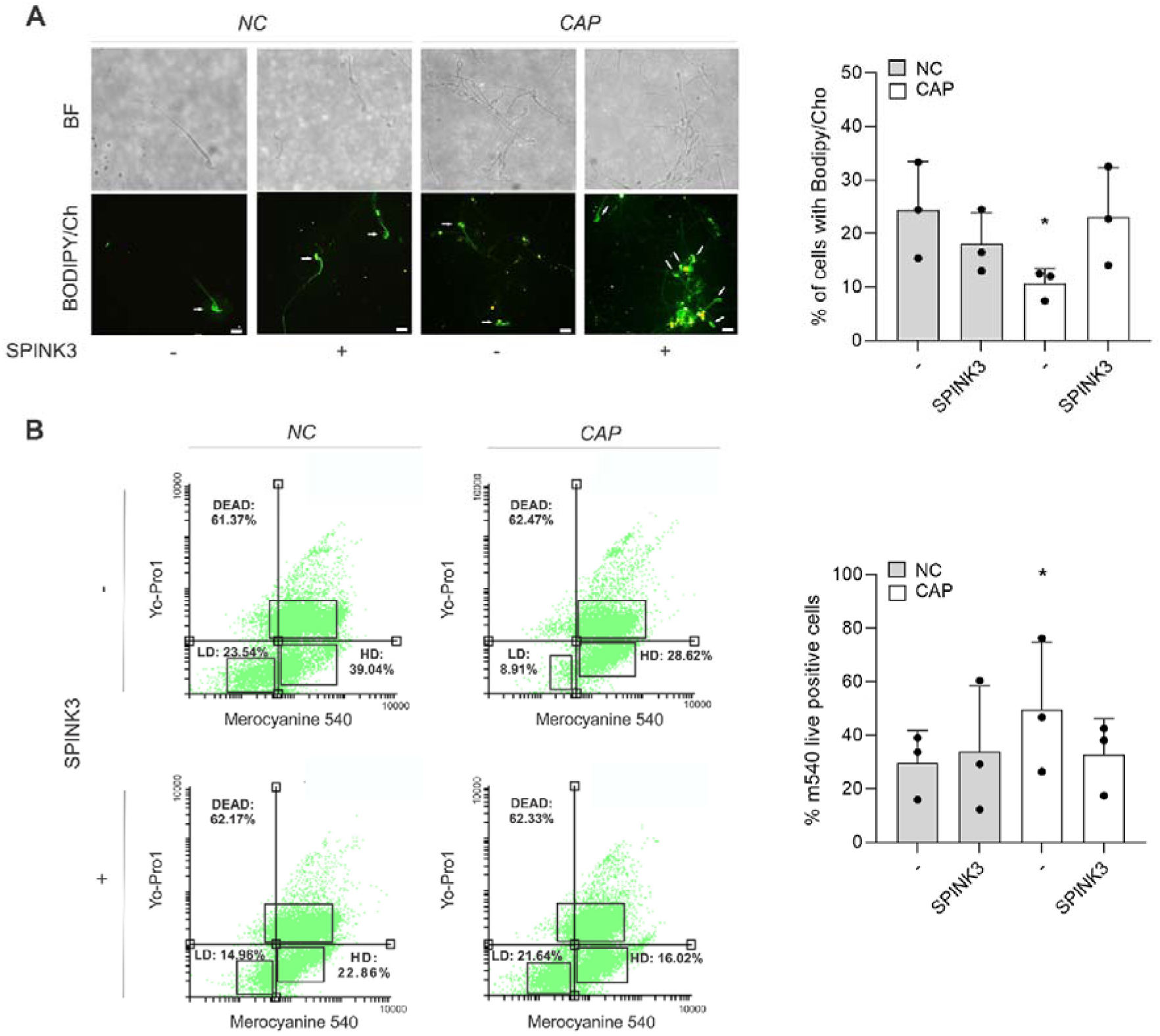
SPINK3 blocks capacitation-associated cholesterol efflux and membrane destabilization. Mouse epididymal spermatozoa were loaded with Bodipy Cholesterol (BODIPY/Ch) **(A)** or M540 and Yo-Pro1 **(B)** with or without SPINK3 (13 µM) and incubated under capacitating (CAP) or non-capacitating (NC) conditions. n=3 LD: low disordered membrane (Yo-Pro1^-^/M540^-^). HD: high disordered membrane (Yo-Pro1^-^/M540^+^). Percentage of M540^+^ cells is expressed relative to the total live cells (Yo-Pro1^-^). (*) Significant differences among treatments, p < 0.05. Scale bar: 10 μm.

### SPINK3-exposed spermatozoa during capacitation reduce *in vitro* fertilization

Pharmacological inhibition of CatSper during *in vitro* capacitation compromises sperm fertilizing ability *in vitro* (Curci et al., 2021). *In vivo*, decapacitation factors are gradually removed from the sperm surface within the female reproductive tract, thereby allowing capacitation to proceed. In contrast, conventional *in vitro* fertilization (IVF) assays are performed with epididymal sperm, which naturally lack seminal plasma proteins such as SPINK3, and membrane-bound proteins might not be efficiently detached under these conditions. To specifically evaluate whether exogenously added SPINK3 exerts an inhibitory effect in vitro, epididymal sperm were incubated under capacitating conditions in the absence or presence of 13 µM recombinant SPINK3 and subsequently used for inseminating cumulus–oocyte complexes (COCs). We found that SPINK3 produced a significant decrease in the percentage of fertilized eggs (**Figure 5A, B**). Most sperm that were preincubated with SPINK3 retained SPINK3 immunostaining in the flagellum when recovered 4 hours after insemination (**Figure 5C**).

**Figure 5.**
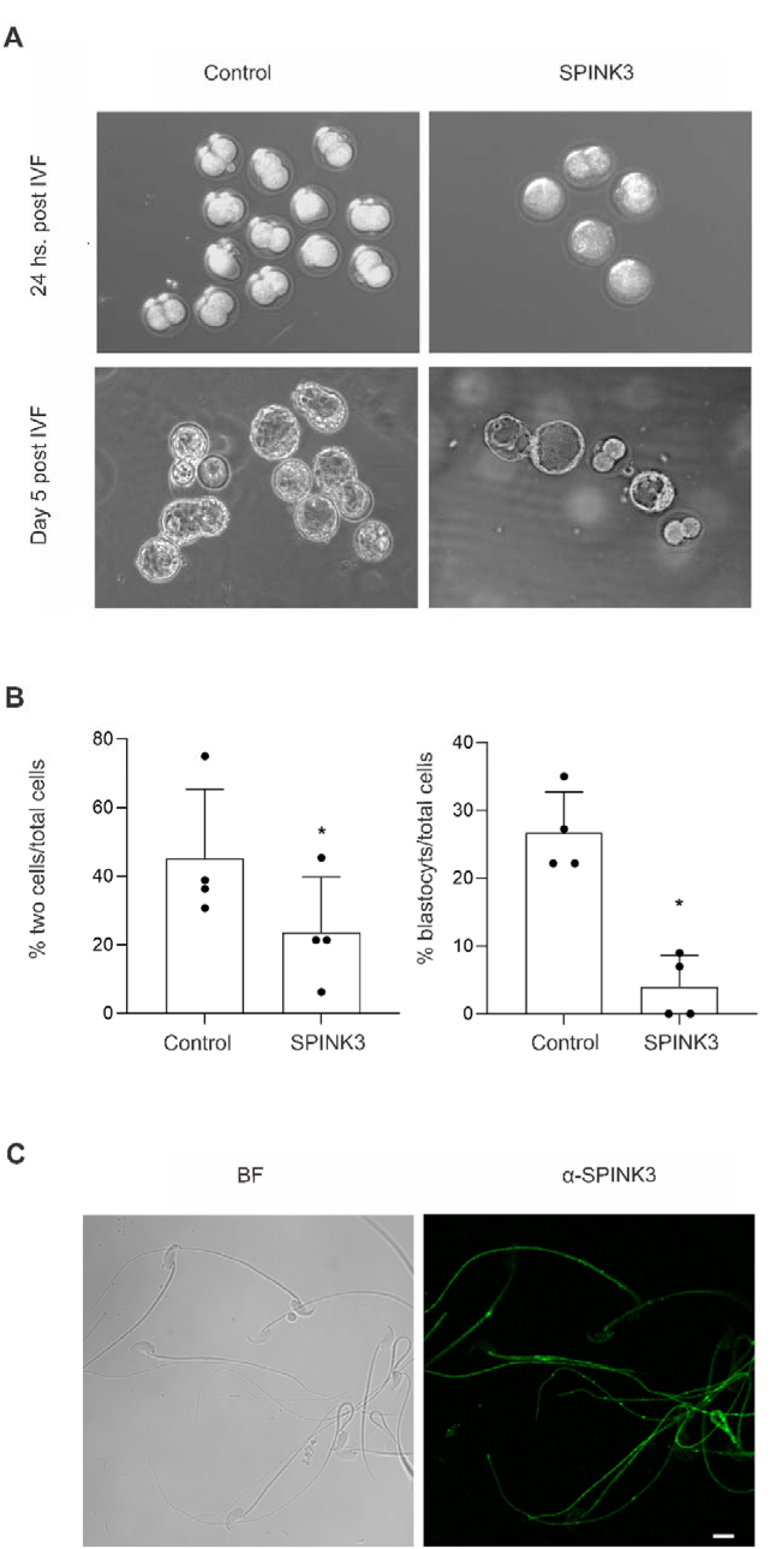
Effect of SPINK3 on *in vitro* fertilization. Mouse sperm were incubated in the absence or presence of rSPINK3 (13 µM) under capacitating conditions (CAP). After capacitation, sperm (5 thousand total cells, 10 µl) were added to groups of COCs (5-6 COCs, 90 µl) obtained from superovulated females, (n=4). **A)** Representative images of two-cell embryos and blastocysts obtained 24 hs. or 5 days post-IVF, respectively. **B)** Quantification of two-cell embryos and blastocysts relative to the total oocytes (n=113). (*) Significant differences compared to the control without SPINK3, p < 0.05. **C)** Sperm recovered after 4 hs. of co-culture were incubated with α-SPINK3 antibodies to detect sperm-bound SPINK3 bound to sperm by immunofluorescence. Scale bar: 10 μm.

## DISCUSSION

### SPINK3 is involved in the regulation of CATSPER1 processing protease in mice

We have previously reported that the small vesicular protein SPINK3, found in mouse male secretions, acts as a decapacitation factor by inhibiting CatSper activity and the increase of [Ca^2+^]_i_ during the early stages of capacitation induced by BSA and HCO ^-^ (Zalazar et al., 2020). We have also shown that SPINK3 functions as a Kazal-type serine protease inhibitor that competitively inhibits trypsin-like proteases (Assis et al., 2013). Concurrently, it has been shown that both CATSPER1 and CATSPERθ undergo capacitation-induced degradation (Chung et al., 2014; Ded et al., 2020; Huang et al., 2023). However, endogenous protein(s) responsible for this physiological regulation remains unidentified. In this study, we show that SPINK3 inhibits sperm trypsin-like proteases and specifically inhibits the capacitation-associated cleavage of CATSPER1, but not CATSPERθ, suggesting a sophisticated regulatory role for SPINK3 in controlling CatSper channel activity.

It has been suggested that inhibitors of calpain, a family of calcium-dependent cysteine proteases, also block CATSPER1 processing (Ded et al., 2020). However, SPINK3, as a serine protease inhibitor, is unlikely to directly inhibit calpains. Given that SPINK3 specifically inhibits serine proteases and interferes with CATSPER1 processing, if this effect results from a direct action of SPINK3 on a sperm protease, then the protease responsible for CATSPER1 cleavage is likely a serine protease. Alternatively, CATSPER1 N-terminal domain cleavage could be a multi-protease event involving different types of proteases. Other sperm serine proteases have been reported in mouse sperm, such as ACROSIN (Q3ZB06_MOUSE), PRS55 (Q14BX2_MOUSE), and TESP1 (Q3U132_MOUSE), but all are localized in the sperm head, either as part of the membrane fraction (GPI-anchored or transmembrane included) or soluble in the acrosome. However, no protease has been described specifically in the sperm flagella, where CATSPER1 processing is thought to occur. This requires further investigation.

### SPINK3 modulates capacitation through mechanisms that extends beyond its protease inhibitory activity

While the finding that SPINK3 inhibits CATSPER1 processing does not fully explain the mechanism by which this protein regulates capacitation, we used soybean trypsin inhibitor (SBTI) as an alternative trypsin-like protease inhibitor to further investigate this process. SBTI had previously been shown to block the SPINK3 binding site on the sperm head in mouse sperm (Ramachandran et al., 2021), suggesting a potential shared target between these molecules in this region. Additionally, SBTI has been used to detect acrosin exposure in other mammalian sperm (Sumigama et al., 2015), indicating its interaction with acrosomal proteases. Given these findings, we selected SBTI to determine whether inhibiting serine proteases alone is sufficient to regulate capacitation. Our results show that, despite SBTI’s ability to inhibit sperm proteases involved in CATSPER1 degradation, it does not modulate capacitation as SPINK3 does, suggesting that SPINK3’s role in capacitation extends beyond its inhibitory function on serine proteases.

Our *in vitro* results demonstrate that both SPINK3 and seminal vesicles extracts (containing native SPINK3) strongly inhibit sperm trypsin-like proteases, however evidence shows that SPINK3 acts extracellularly. Therefore, it is possible to consider a role for surface attached SPINK3 in stabilizing the membrane and CatSper arrangements in the principal piece, by binding a membrane protease, whose substrate is CATSPER1, and likely to be differently compartmentalized outside of CatSper nanodomains before capacitation. Although there is no strict co-localization of SPINK3 and CATSPER1, sperm from *Catsper1* KO mice, which lack the entire channel complex in the spermatozoa, displayed a different SPINK3-sperm binding ability and pattern, suggesting that SPINK3 binds to the sperm flagellum in a CatSper dependent manner. The different and compromised SPINK3 binding with sperm may be because lipid domain arrangement was disrupted due to the lack of CatSper channel in the sperm flagella. Given that TESP1 has previously been reported as the ligand for SPINK3 in the acrosomal region (Ramachandran et al., 2021), it is possible that the mechanisms governing SPINK3 binding to sperm vary across membrane subregions. In sperm lacking the CatSper channel—and consequently exhibiting defective molecular organization along the flagellum—SPINK3 attachment may be restricted to the head, as the principal piece may no longer provide a suitable binding site. Additionally, SPINK3 might play distinct roles in different sperm regions: one regulating calcium influx to modulate hyperactivation and the other controlling the acrosome reaction. However, the latter is beyond the scope of this study and requires further investigation.

### SPINK3 might be involved in the development of sperm subpopulations in the oviduct

SPINK3 is bound to the majority of sperm during ejaculation, and it is progressively detached along the oviduct; however, this detachment is asynchronous, as it does not occur simultaneously in all sperm (Nicolli et al., 2022). Besides, it is not clear what modulates CatSper activity and how CatSper channel containing N-terminally processed CATSPER1 (Ded et al., 2020) alters its activity. As most sperm found in the ampulla has a molecular barcode of acrosome reacted (AR +), tyrosine phosphorylation negative (pY -), and CATSPER1 intact (CATSPER1 +), and our results show that pY-sperm are also SPINK3+, one can speculate that late release of SPINK3 from the flagellar membrane contributes to avoid premature cleavage of CATSPER1 and thus premature degeneration. Proteolysis of membrane protein ectodomains acts as a post-translational modification that determines the levels and activity of membrane proteins (Lichtenthaler et al., 2018). Would CATSPER1-cleaved sperm show increased or reduced CatSper activity? Would intact CATSPER1 be maintained in the fertilizing sperm or just until right before? In this work, our evidence shows that SPINK3 prevents this proteolysis that occurs in capacitating media and also prevents membrane reorganization, increase in [Ca^2+^]_i_, and acrosome exocytosis, but it is unclear whether early or late release constitutes the destiny of sperm for fertilization. Recently, it has been reported that seminal plasma spermine reversibly inhibits CatSper’s temperature gating, protecting against premature activation. May heterogeneity reflects a compilation of the time-and space-dependent changes that sperm undergo along the oviduct? Could the importance of tightly regulating CatSper function be such that multiple redundant molecules operate in parallel to ensure fertilization success? All these questions serve ongoing and future research directions.

While SPINK3 is gradually removed from sperm surface *in vivo* in female’s reproductive tract (Nicolli et al., 2022), the detachment is less efficient *in vitro*, as evidenced by the intense SPINK3 labeling in the sperm recovered from the co-culture and fertility significantly reduced, suggesting that the detachment of SPINK3 from sperm needs specific physiological conditions or specific molecules in female’s reproductive tract.

In conclusion, we demonstrate a role for SPINK3 in stabilizing the membrane and preventing CATSPER1 processing, either indirectly or directly acting as an inhibitor of the responsible protease that cleaves CATSPER1. Our findings provide molecular insights into the dynamic regulation of capacitation by male-derived SPINK3 and provide the first evidence of a natural molecule involved in modulating the CatSper channel. This work helps to explain the mechanism for the graded subpopulations of sperm along the female reproductive tract.

## Acknowledgements

The authors would like to thank Viviana Daniel for their assistance during confocal microscopy. The authors also acknowledge Hugo Nuñez for his assistance with the animal handling and maintenance.

## Data availability statement

All datasets used for this study are available upon request.

## Funding

This work was supported by the National Scientific and Technical Research Council (CONICET, Argentina, PIP 11220200100850CO) awarded to A.C., the National Institutes of Health (R01HD096745) to J.-J.C, the Chan Zuckerberg Initiative (2021-240504), the National Institutes of Health (R01HD106968), and the Agencia Nacional de Promoción de la Investigación, el Desarrollo Tecnológico y la Innovación (Argentina, 2020-00988) to M.G.B, and grants PICT 2019-1779 and PICT 2021-0102 to D.K.. X.H. is a postdoctoral fellowship awardee of Male Contraceptive Initiative (MCI).

### Author contribution statement

AC and JJ contributed to the conceptualization of the study and supervision. AN, XH, CS, GD, AR and LG contributed to the design and development of the methodology. AN, XH, DK, MB, LZ, AC and JJ contributed to the writing and editing of the manuscript. All the co-authors were involved in the formal analysis, validation, visualization of the results and revised the manuscript.

---

**Supplementary Figure 1.**
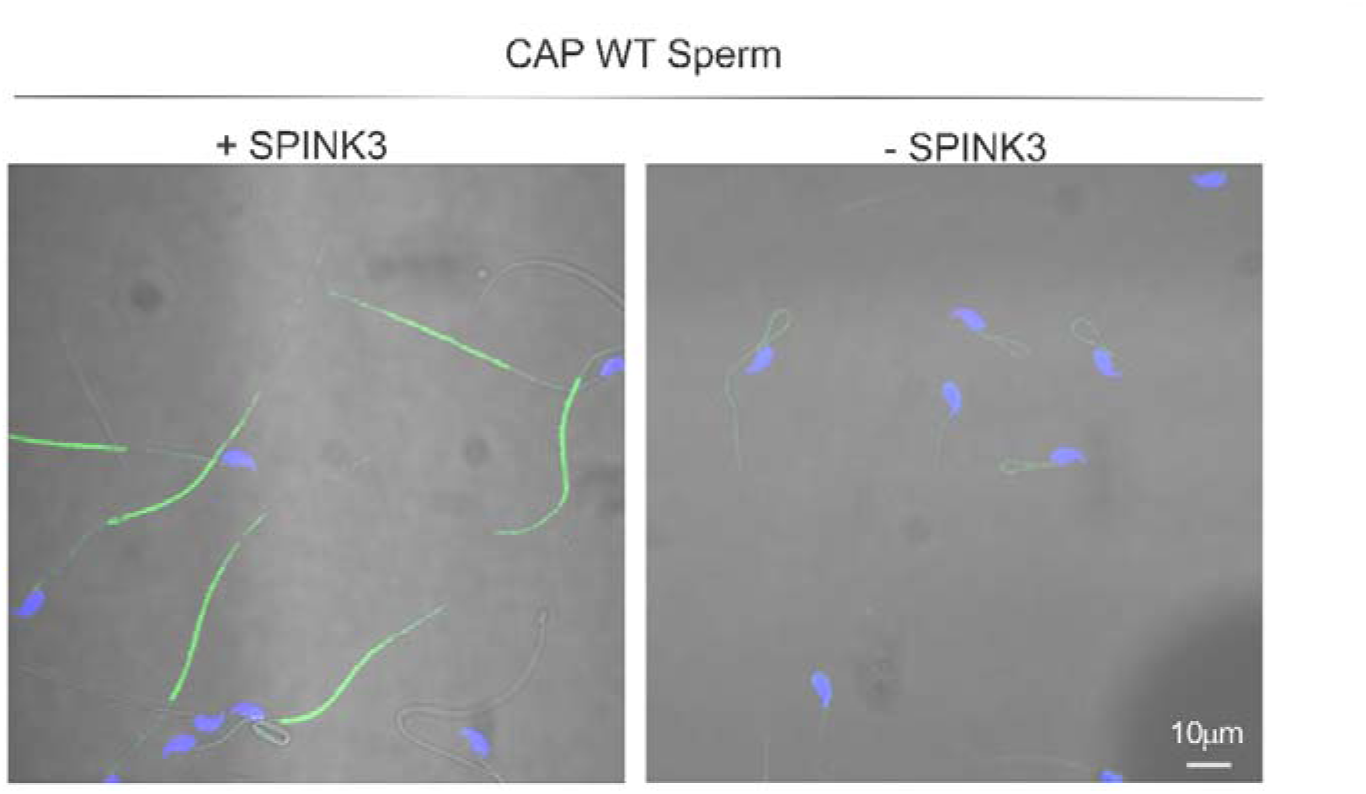
Immunostaining was performed with anti-CATSPER1 antibody on capacitated sperm that were previously incubated w/o SPINK3 for 30 min. Representative images from three independent experiments are shown. Scale bar: 10 μm.

**Supplementary Figure 2.**
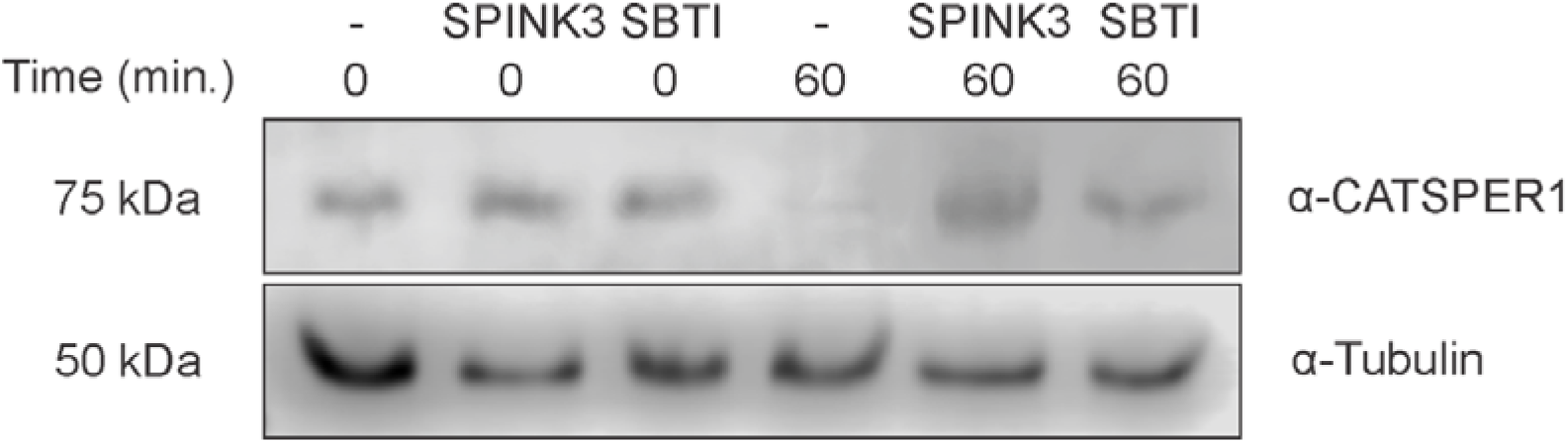
*Effect of SPINK3 and SBTI on the proteolytic degradation of CATSPER1 in vivo.* Mouse epididymal sperm (n=3) were incubated with or without SPINK3 (13 µM) or SBTI (1 µM) and exposed to CAP conditions. Cell lysis was performed on aliquots of different treatments at 0 and 60 min. Proteins (0.5 x 10^6^ cells/ml) were separated on a 10% polyacrylamide gel and revealed by Western blot, using anti-CATSPER1 followed by anti-Tubulin.

**Supplementary Figure 3.**
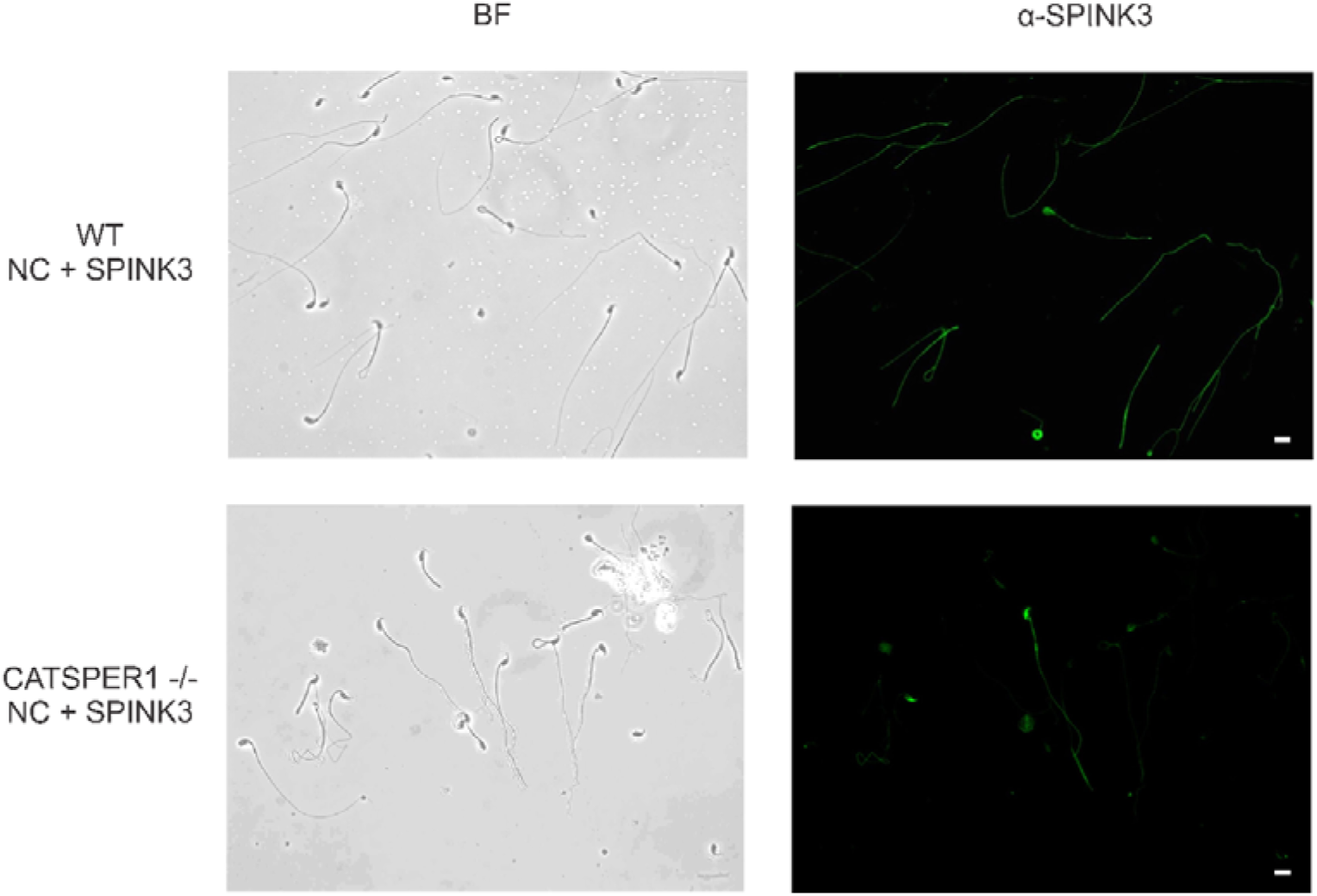
*Immunodetection of recombinant SPINK3 incubated with epididymal spermatozoa from wild type and Catsper1 KO mice.* Mouse epididymal NC spermatozoa were incubated in the presence of SPINK3 (13 μM) and immunofluorescence assays were performed using rabbit anti-SPINK3 (α-SPINK3) followed by anti-IgG-rabbit-Alexa488 (green). (n=3) Scale bar: 10 μm. Images are representative of three independent experiments. BF, bright field.

**Supplementary Movie 1.** *3D projection of SPINK3 and plasma membrane markers FM4-64 FX and CellBrite Fix on the sperm flagellum, related to* Figure 2. The video shows a rotating view along the y-axis, providing a spatial visualization of SPINK3 (red) localization on the plasma membrane of the sperm principal piece. The membrane is labeled with FM4-64 FX (green) and CellBrite Fix (cyan), highlighting differences in staining intensity between the midpiece and the principal piece. https://drive.google.com/drive/folders/1nl2lqlkbBCWcofPsq-asTwwj_lwKv36C?usp=drive_link

## REFERENCES

1. Assis, D. M., Zalazar, L., Magno, D., 1#, A., Juliano, M. A., De Castro, R., & Cesari, A. (2013). Novel inhibitory activity for serine protease inhibitor Kazal type-3 (Spink3) on human recombinant kallikreins. Ingentaconnect.Com, 20, 1098–1107. 10.2174/0929866511320100003

2. Austin, C. R. (1952). The capacitation of the mammalian sperm. Nature, 170(4321). 10.1038/170326a0

3. Bernecic, N. C., Zhang, M., Gadella, B. M., Brouwers, J. F. H. M., Jansen, J. W. A., Arkesteijn, G. J. A., de Graaf, S. P., & Leahy, T. (2019). BODIPY-cholesterol can be reliably used to monitor cholesterol efflux from capacitating mammalian spermatozoa. Scientific Reports, 9(1). 10.1038/s41598-019-45831-7

4. Cesari, A., Monclus, aM. de los A., Tejón, G. P., Clementi, M., & Fornes, M. W. (2010). Regulated serine proteinase lytic system on mammalian sperm surface: There must be a role. Theriogenology, 74(5), 699–711.e5. 10.1016/J.THERIOGENOLOGY.2010.03.029

5. Chung, J. J., Navarro, B., Krapivinsky, G., Krapivinsky, L., & Clapham, D. E. (2011). A novel gene required for male fertility and functional CATSPER channel formation in spermatozoa. Nature Communications, 2(1). 10.1038/ncomms1153

6. Chung, J. J., Shim, S. H., Everley, R. A., Gygi, S. P., Zhuang, X., & Clapham, D. E. (2014). Structurally distinct Ca2+ signaling domains of sperm flagella orchestrate tyrosine phosphorylation and motility. Cell, 157(4). 10.1016/j.cell.2014.02.056

7. Curci, L., Carvajal, G., Sulzyk, V., Gonzalez, S. N., & Cuasnicú, P. S. (2021). Pharmacological Inactivation of CatSper Blocks Sperm Fertilizing Ability Independently of the Capacitation Status of the Cells: Implications for Non-hormonal Contraception. Frontiers in Cell and Developmental Biology, 9. 10.3389/fcell.2021.686461

8. Ded, L., Hwang, J. Y., Miki, K., Shi, H. F., & Chung, J. J. (2020). 3D in situ imaging of female reproductive tract reveals molecular signatures of fertilizing spermatozoa in mice. ELife, 9. 10.7554/eLife.62043

9. Dematteis, A., Miranda, S., … M. N.-B. of, & 2008, undefined. (n.d.). Rat caltrin protein modulates the acrosomal exocytosis during sperm capacitation. Academic.Oup.Com. Retrieved April 2, 2023, from https://academic.oup.com/biolreprod/article-abstract/79/3/493/2557586

10. Dukelow, W. R., Chernoff, H. N., & Williams, W. L. (1967). Properties of decapacitation factor and presence in various species. Journal of Reproduction and Fertility, 14(3). 10.1530/jrf.0.0140393

11. Ferreira, J. J., Lybaert, P., Puga-Molina, L. C., & Santi, C. M. (2021). Conserved Mechanism of Bicarbonate-Induced Sensitization of CatSper Channels in Human and Mouse Sperm. Frontiers in Cell and Developmental Biology, 9. 10.3389/fcell.2021.733653

12. Flesch, F. M., Brouwers, J. F. H. M., Nievelstein, P. F. E. M., Verkleij, A. J., Van Golde, L. M. G., Colenbrander, B., & Gadella, B. M. (2001). Bicarbonate stimulated phospholipid scrambling induces cholesterol redistribution and enables cholestrol depletion in the sperm plasma membrane. Journal of Cell Science, 114(19). 10.1242/jcs.114.19.3543

13. Huang, X., Miyata, H., Wang, H., Mori, G., Iida-Norita, R., Ikawa, M., Percudani, R., & Chung, J. J. (2023). A CUG-initiated CATSPERθ functions in the CatSper channel assembly and serves as a checkpoint for flagellar trafficking. Proceedings of the National Academy of Sciences of the United States of America, 120(39). 10.1073/pnas.2304409120

14. Hwang, J. Y. (2024). Analysis of Ca2+-mediated sperm motility to evaluate the functional normality of the sperm-specific Ca2+ channel, CatSper. Frontiers in Cell and Developmental Biology, 12. 10.3389/fcell.2024.1284988

15. Hwang, J. Y., & Chung, J. J. (2023). CatSper Calcium Channels: 20 Years On. In Physiology (Vol. 38, Issue 3). 10.1152/PHYSIOL.00028.2022

16. Hwang, J. Y., Mannowetz, N., Zhang, Y., Everley, R. A., Gygi, S. P., Bewersdorf, J., Lishko, P. V., & Chung, J. J. (2019). Dual Sensing of Physiologic pH and Calcium by EFCAB9 Regulates Sperm Motility. Cell, 177(6). 10.1016/j.cell.2019.03.047

17. Kirichok, Y., Navarro, B., & Clapham, D. E. (2006). Whole-cell patch-clamp measurements of spermatozoa reveal an alkaline-activated Ca2+ channel. Nature, 439(7077). 10.1038/nature04417

18. Leahy, T., & Gadella, B. M. (2011). Sperm surface changes and physiological consequences induced by sperm handling and storage. In Reproduction (Vol. 142, Issue 6). 10.1530/REP-11-0310

19. Lenth, R. V. (2016). Least-squares means: The R package lsmeans. Journal of Statistical Software, 69. 10.18637/jss.v069.i01

20. Li, J., Roca, J., Pérez-Patiño, C., … I. B.-A. reproduction, & 2018, undefined. (n.d.). Is boar sperm freezability more intrinsically linked to spermatozoa than to the surrounding seminal plasma? Elsevier. Retrieved April 2, 2023, from https://www.sciencedirect.com/science/article/pii/S0378432017309363

21. Lichtenthaler, S. F., Lemberg, M. K., & Fluhrer, R. (2018). Proteolytic ectodomain shedding of membrane proteins in mammals—hardware, concepts, and recent developments. The EMBO Journal, 37(15). 10.15252/embj.201899456

22. Lin, H., Luo, C., Wang, C., Biophysical, Y. C.-B. and, & 2006, undefined. (n.d.). Epitope topology and removal of mouse acrosomal plasma membrane by P12-targeted immunoaggregation. Elsevier. Retrieved April 2, 2023, from https://www.sciencedirect.com/science/article/pii/S0006291X06018377

23. Lin, M., Lee, R., Hwu, Y., Lu, C., … S. C.-, & 2008, undefined. (n.d.). SPINKL, a Kazal-type serine protease inhibitor-like protein purified from mouse seminal vesicle fluid, is able to inhibit sperm capacitation. Rep.Bioscientifica.Com. Retrieved April 2, 2023, from https://rep.bioscientifica.com/view/journals/rep/136/5/559.xml

24. McEvoy, L., Schlegel, R. A., Williamson, P., & Del Buono, B. J. (1988). Merocyanine 540 as a flow cytometric probe of membrane lipid organization in leukocytes. Journal of Leukocyte Biology, 44(5). 10.1002/jlb.44.5.337

25. Nature, M. C.-, & 1951, undefined. (n.d.). Fertilizing capacity of spermatozoa deposited into the fallopian tubes. Springer. Retrieved April 2, 2023, from https://link.springer.com/content/pdf/10.1038/168697b0.pdf

26. Nicolli, A., Alonso, C., Otamendi, C., … M. C.-, & 2022, undefined. (n.d.). How, where and when is SPINK3 bound and removed from mouse sperm? *Rep.Bioscientifica.Com*. Retrieved April 2, 2023, from https://rep.bioscientifica.com/view/journals/rep/163/5/REP-21-0195.xml

27. Nicolli, A. R., & Cesari, A. (2023). The other side of capacitation: role of mouse male molecules in the regulation of time and place of capacitation. Reproduction, 166(6). 10.1530/REP-23-0188

28. Noda, T., & Ikawa, M. (2019). Physiological function of seminal vesicle secretions on male fecundity. In Reproductive Medicine and Biology (Vol. 18, Issue 3). 10.1002/rmb2.12282

29. Pinheiro, J., Bates, D., DebRoy, S., Sarkar, D., Heisterkamp, S., & Van Willigen, B. (2020). Package nlme: Linear and Nonlinear Mixed Effects Models. *R Package*.

30. Qi, H., Moran, M. M., Navarro, B., Chong, J. A., Krapivinsky, G., Krapivinsky, L., Kirichok, Y., Ramsey, I. S., Quill, T. A., & Clapham, D. E. (2007). All four CatSper ion channel proteins are required for male fertility and sperm cell hyperactivated motility. Proceedings of the National Academy of Sciences of the United States of America, 104(4). 10.1073/pnas.0610286104

31. Quill, T. A., Sugden, S. A., Rossi, K. L., Doolittle, L. K., Hammer, R. E., & Garbers, D. L. (2003). Hyperactivated sperm motility driven by CatSper2 is required for fertilization. Proceedings of the National Academy of Sciences of the United States of America, 100(25). 10.1073/pnas.2136654100

32. Ramachandran, S., … R. B.-M. H., & 2021, undefined. (n.d.). A mouse testis serine protease, TESP1, as the potential SPINK3 receptor protein on mouse sperm acrosome. Academic.Oup.Com. Retrieved April 2, 2023, from https://academic.oup.com/molehr/article-abstract/27/10/gaab059/6370709

33. Ren, D., Navarro, B., Perez, G., Jackson, A. C., Hsu, S., Shi, Q., Tilly, J. L., & Clapham, D. E. (2001). A sperm ion channel required for sperm motility and male fertility. Nature, 413(6856). 10.1038/35098027

34. Samanta, L., Parida, R., Dias, T. R., & Agarwal, A. (2018). The enigmatic seminal plasma: A proteomics insight from ejaculation to fertilization. In Reproductive Biology and Endocrinology (Vol. 16, Issue 1). 10.1186/s12958-018-0358-6

35. Sankaranarayanan, S., Kellner-Weibel, G., De La Llera-Moya, M., Phillips, M. C., Asztalos, B. F., Bittman, R., & Rothblat, G. H. (2011). A sensitive assay for ABCA1-mediated cholesterol efflux using BODIPY-cholesterol. Journal of Lipid Research, 52(12). 10.1194/jlr.D018051

36. Silva, A. A. S., Raimundo, T. R. F., Mariani, N. A. P., Kushima, H., Avellar, M. C. W., Buffone, M. G., Paula-Lopes, F. F., Moura, M. T., & Silva, E. J. R. (2021). Dissecting EPPIN protease inhibitor domains in sperm motility and fertilizing ability: Repercussions for male contraceptive development. Molecular Human Reproduction, 27(12). 10.1093/molehr/gaab066

37. Steckler, D., Stout, T. A. E., Durandt, C., & Nöthling, J. O. (2015). Validation of merocyanine 540 staining as a technique for assessing capacitation-related membrane destabilization of fresh dog sperm. Theriogenology, 83(9). 10.1016/j.theriogenology.2015.01.019

38. Sumigama, S., Mansell, S., Miller, M., Lishko, P. V., Cherr, G. N., Meyers, S. A., & Tollner, T. (2015). Progesterone accelerates the completion of sperm capacitation and activates catsper channel in spermatozoa from the rhesus macaque. Biology of Reproduction, 93(6). 10.1095/biolreprod.115.129783

39. Tiwari, R., Manzar, N., Bhatia, V., Yadav, A., Nengroo, M. A., Datta, D., Carskadon, S., Gupta, N., Sigouros, M., Khani, F., Poutanen, M., Zoubeidi, A., Beltran, H., Palanisamy, N., & Ateeq, B. (2020). Androgen deprivation upregulates SPINK1 expression and potentiates cellular plasticity in prostate cancer. Nature Communications, 11(1). 10.1038/s41467-019-14184-0

40. Torres-García, E., Pinto-Cámara, R., Linares, A., Martínez, D., Abonza, V., Brito-Alarcón, E., Calcines-Cruz, C., Valdés-Galindo, G., Torres, D., Jabloñski, M., Torres-Martínez, H. H., Martínez, J. L., Hernández, H. O., Ocelotl-Oviedo, J. P., Garcés, Y., Barchi, M., D’Antuono, R., Bošković, A., Dubrovsky, J. G., … Guerrero, A. (2022). Extending resolution within a single imaging frame. Nature Communications, 13(1). 10.1038/s41467-022-34693-9

41. Zalazar, L., Stival, C., Nicolli, A. R., De Blas, G. A., Krapf, D., & Cesari, A. (2020). Male Decapacitation Factor SPINK3 Blocks Membrane Hyperpolarization and Calcium Entry in Mouse Sperm. Frontiers in Cell and Developmental Biology, 8. 10.3389/FCELL.2020.575126/FULL

42. Zuur. (2019). Mixed Effects Models and Extensions in Ecology with R by Zuur. In springer (Vol. 53, Issue 9).

