## Supplementary Information S1 for "SPINK3-sperm interaction determines a stable sperm subpopulation with intact CatSper channel"

### Recombinant SPINK3 production and characterization

#### *Cloning and expression of recombinant SPINK3*

A DNA fragment encoding the mature form of mouse SPINK3 (residues 24 to the stop codon, nucleotides 158–331) was cloned in the expression vector pET-24b (+) (Novagen) as previously described (19). Briefly, the cDNA corresponding to mature SPINK3 was PCR-amplified from pGEX-SPINK3 and subcloned into the pET-24b(+) vector (Novagen). NdeI and XhoI were added to the 5' end nucleotide sequences of Spink3-Fw (5'-GG GAA TTC CAT ATG GCT AAG GTG ACT GG-3') and Spink3-Rev (5'-GGC CTC GAG GCA AGG CCC ACC-TTT TC-3') primers, respectively. The reverse primer excluded the stop codon to allow in-frame fusion to a C-terminal $His_6$-tag. The construct was verified by DNA sequencing and transformed into *E. coli* Rosetta™ (DE3) cells. For overexpression of Spink3-His6, *E. coli* Rosetta cells (Novagen) were transformed with the pET-24b (+)- SPINK3 and the cells harboring the plasmid were grown overnight at 37 °C in Luria Bertani (2YT-LB) medium (1% w/v yeast extract, 2% w/v tryptone, 0.5% w/v NaCl) supplemented with kanamycin (50 g/ml) and chloramphenicol (35 g/ml). This starting culture was used to inoculate (1/100) the same medium and the culture was grown at 37 °C under continuous shaking until the optical density at 600 nm (OD600) reached 0.4, then shifted to 18 °C and further incubated to an OD600 ~0.6. Synthesis of SPINK3 was induced by addition of 0.1 mM IPTG for 16 h at 18° C.

The recombinant design of SPINK3, previously established to express the mature form of the protein, takes into account its eukaryotic origin and allows efficient expression in E. coli, a system that supports disulfide bond formation and solubility. According to UniProt (P09036·ISK1_MOUSE), SPINK3 does not present experimentally validated glycosylation. SignalP predictions indicate that the mature native protein begins at residue 24, and our recombinant design lacks the signal peptide, ensuring expression of the mature form. NetNGlyc 1.0 and NetOGlyc 4.0.0.13 analyses predict no N-linked or O-linked glycosylation sites, respectively. Therefore, no post-translational modifications are expected in the recombinant protein expressed in *E. coli*, as further supported by its consistent migration in SDS-PAGE and functional activity.

#### *Cell lysis and clarification*

Bacterial pellets were collected by centrifugation (1700 × g, 8 min, 4 °C), washed twice with PBS 1×, and resuspended in lysis buffer (20 mM phosphate buffer pH 7.4, 137 mM NaCl, 2.7 mM KCl). Lysozyme (1 mg/ml final, 15 mg total) was added, and suspensions were frozen at –80 °C overnight. After thawing on ice, 0.2% Triton X-100 was added. Lysis was performed by sonication (6 cycles × 30 s on ice, 40 W). Lysates were clarified by centrifugation (12,000 × g, 30 min, 4 °C). The soluble fraction (sperm lysate) was collected and kept on ice.

#### *Purification of SPINK3-$His_6$*

The clarified lysate was applied to a HiTrap IMAC HP column (GE Healthcare) pre-charged with 0.1 M $NiSO_4$ and equilibrated in binding buffer (20 mM sodium phosphate, 0.5 M NaCl, 30 mM imidazole, pH 7.4). Recombinant SPINK3 was eluted in three sequential steps using elution buffers containing increasing concentrations of imidazole: 120 mM, 200 mM, and 500 mM. The collected fractions were subsequently analyzed by SDS-PAGE as described below. Fractions with similar protein concentration and purity were pooled and dialyzed against PBS.

#### *SDS-PAGE and western blot*

Aliquots of soluble lysate and purified protein were mixed with sample buffer (2 µl) and DTT (1 µl), heated 5 min at 95 °C, and separated by 15% SDS-PAGE. Gels were stained with Coomassie Brilliant Blue to assess purity (**Fig. SI 1A**). For immunoblotting, proteins were transferred onto PVDF membranes (2 h, 80 mA) in Towbin buffer. Membranes were blocked in 3% BSA in TBST and incubated overnight with anti-SPINK1 antibody (1:2500, Sigma HPA027498). After washing, membranes were incubated with alkaline phosphatase-conjugated anti-rabbit IgG-HRP conjugated (1:10000, A615Y Sigma-Aldrich) and revealed using a chemiluminescence detection kit (ECL plus, Amersham, GE Healthcare) as per the manufacturer's instructions, and signal was detected with a Licor C-Digit chemiluminescence scanner (**Fig. SI 1B**).

#### *Trypsin activity assay*

The inhibitory activity of recombinant SPINK3 was measured using the fluorogenic substrate Z-Arg-Gly-Arg-AMC (CGGR-AMC, Sigma). All reactions were performed in 96-well black plates (Nunc) with a final volume of 100 µl per well.

Reaction mixtures prepared at room temperature and included Trypsin (2.7 nM final), 50 mM Buffer phosphate pH 7.4, w/o the following inhibitors: SPINK3 (1.3 µM final), SBTI (2 µg/ml final), seminal vesicle homogenate as a control of native SPINK3 (6 µl). The reaction was initiated by adding 2 µM CGGR-AMC substrate, and fluorescence (Ex 355 nm / Em 436 nm) was measured every minute for 30 min at 37 °C with agitation (Fluoroskan Ascent microplate reader, Thermo Electron Corporation, Waltham, MA, USA). Enzymatic activity was calculated as Δfluorescence over time and expressed relative to the trypsin control. Inhibition is evidenced as a decrease in proteolytic activity (**Fig. SI 1C**).

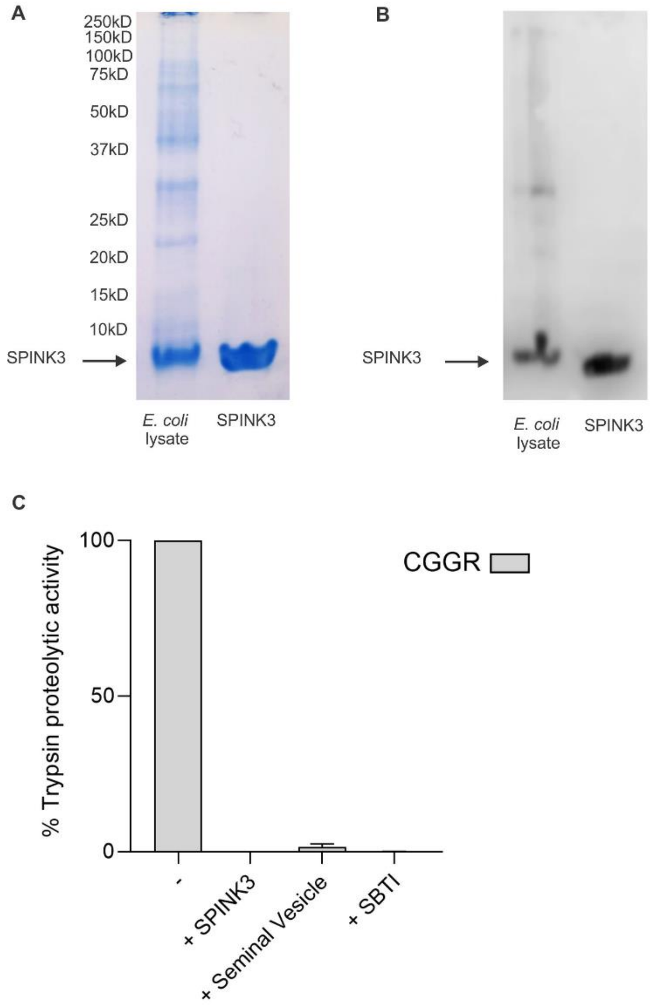


**Figure SI1. Expression, detection, and inhibitory activity of recombinant SPINK3. A)** Coomassie-stained SDS-PAGE of *E. coli* lysate showing the overexpression of SPINK3 and purified SPINK3. The arrow indicates the expected band corresponding to recombinant protein. **B)** Western blot with anti-SPINK1 antibodies. **C)** Proteolytic activity assay over the specific trypsin substrate (CGGR). Trypsin activity is inhibited in the presence of recombinant SPINK3, seminal vesicle extract, or SBTI (soybean trypsin inhibitor), demonstrating the inhibitory capacity of recombinant SPINK3.
